# Spatial and ecological drivers of population structure in *Alkanna tinctoria* (Boraginaceae), a polyploid medicinal herb

**DOI:** 10.1101/2021.05.06.442843

**Authors:** Muhammad Ahmad, Thibault Leroy, Nikos Krigas, Eva M. Temsch, Hanna Weiss-Schneeweiss, Christian Lexer, Eva Maria Sehr, Ovidiu Paun

**Affiliations:** Center for Health & Bioresources, AIT Austrian Institute of Technology GmbH, Tulln, Austria; Department of Botany and Biodiversity Research, University of Vienna, Austria; Institute of Plant Breeding and Genetic Resources, Hellenic Agricultural Organization–Demeter (HAO□Demeter), Thessaloniki, Greece

**Author notes:** Shared last authorship (Eva Maria Sehr, Ovidiu Paun). Author of correspondence: Eva Maria Sehr. Deceased (Christian Lexer).

**Keywords:** Alkanna, EBG, genetic variation, Greece, RAD-seq, polyploid

## Abstract

**Background and Aims:** Quantifying genetic variation is fundamental to understand a species’ demographic trajectory and its ability to adapt to future changes. In comparison to diploids, however, genetic variation and factors fostering genetic divergence remains poorly studied in polyploids due to analytical challenges. Here, by employing a ploidy-aware framework, we investigated the genetic structure and its determinants in polyploid *Alkanna tinctoria* (Boraginaceae), an ancient medicinal herb that is the source of bioactive compounds known as alkannin and shikonin (A/S). From a practical perspective, such investigation can inform biodiversity management strategies.

**Methods:** We collected 14 populations of *A. tinctoria* within its main distribution range in Greece and genotyped them using restriction site-associated DNA sequencing (RAD-seq). As an outgroup, we included two populations of *A. sieberi*. By using a ploidy-aware genotype calling based on likelihoods, we generated a dataset of 16,107 high quality SNPs. Classical and model-based analysis was done to characterize the genetic structure within and between the sampled populations. Finally, to reveal the drivers of genetic structure, we searched for associations between allele frequencies and spatial and climatic variables.

**Key Results:** We found support for a marked regional structure in *A. tinctoria* along a latitudinal gradient in line with phytogeographic divisions. Several analyses identified interspecific admixture affecting both mainland and island populations. Modelling of spatial and climatic variables further demonstrated a larger contribution of neutral processes and a lesser albeit significant role of selection in shaping the observed genetic structure in *A. tinctoria*.

**Conclusions:** Current findings provide evidence of strong genetic structure in *A. tinctoria* mainly driven by neutral processes. The revealed natural genomic variation in Greek *Alkanna* can be used to further predict variation in A/S production, whereas our bioinformatics approach should prove useful for the study of other non-model polyploid species.

## 1. INTRODUCTION

Polyploids are very frequent in flowering plants as at least 35% of extant species are of recent polyploid origin (Wood *et al*., 2009). Despite their high abundance in nature, economic, asthetic and medicinal importance, polyploids are still rarely investigated in population genomics (van de Peer *et al*., 2017). One of the reasons is that such investigations remain challenging, due to uncertainty in estimating allele copy number (*i.e*., dosage) and, therefore, to accurately assess genotype and allele frequencies (Meirmans and van Tienderen, 2013; Dufresne *et al*., 2014; Meirmans *et al*., 2018). In addition, the genome complexity of polyploids is often exacerbated by heterogeneous inheritance patterns across chromosomes, ranging from fully tetrasomic to disomic inheritance (Dufresne *et al*., 2014; Meirmans *et al*., 2018).

Genotype likelihood-based methods, (as implemented *e.g.*, in ANGSD; Nielsen *et al*., 2011; Korneliussen *et al*., 2014) that are accounting for uncertainties regarding genotype calling, have been shown to greatly improve accuracy of population genetics parameters estimated from low to medium coverage NGS data in diploid species (Warmuth and Ellegren, 2019). Recently, an accurate Bayesian method for estimating polyploid genotypes at biallelic single nucleotide polymorphisms (SNPs) based on likelihoods has been developed (Blischak *et al*., 2016, 2018), but has not yet been widely applied (but see *e.g*., Brandrud *et al*., 2020). Restriction site-associated DNA sequencing (RAD-seq; Baird *et al*., 2008) is a popular method to generate NGS data and investigate genetic variation and population structure, especially for species not benefiting from an available suitable reference genome (Rochette and Catchen, 2017). However, only a limited number of RAD-seq studies have been employed to investigate genetic variation in non-model polyploid species (Zohren *et al*., 2016; Brandrud *et al*., 2017; Záveská *et al*., 2019; Wagner *et al*., 2020; Wang *et al*., 2021).

Genetic variation is a fundamental determinant of the capacity of populations, species and ecosystems to persist and adapt in the face of environmental change (Hughes *et al*., 2008). More than a single gene pool often exists across a species’ range within a spatial and/or ecological arrangement (Hartl and Clark, 1997; Jones and Wang, 2012). The intensity of gene flow between populations, together with the potential for divergent selection and neutral processes, including drift, are largely defining the degree of population genetic structure (Papadopulos *et al*., 2014). Geographically limited dispersal among individuals produces a spatial genetic structure called isolation by distance (IBD; Vekemans and Hardy, 2004). IBD is quantified as the strength of correlation between genetic dissimilarity and geographic distance (Wright, 1943). However, an increased divergence between populations can be also observed as a result of ecologically-based divergent selection against maladapted migrants and hybrids, which in turn, reduces effective gene flow (Nosil *et al*., 2009), commonly referred as ecologically driven isolation (IBE; Wang and Bradburd, 2014). The distribution of intraspecific genetic variation often reflects geographical proximity, as well as the spatial heterogeneity in the ecological landscape suggesting that functional traits may show a similar pattern, mirroring selective forces (Fournier-Level *et al*., 2011). Hence, investigating the genetic structure of populations together with its drivers is key for understanding the evolutionary history of a group, and predicting its future, with important implications for biodiversity management and conservation.

In the current study, we used a ploidy-aware likelihood-based genotyping to quantify genetic diversity and to investigate the population structure of the polyploid plant *Alkanna tinctoria* Tausch (Boraginaceae) across most of its distribution area in mainland Greece. In addition, as a potential outgroup species, we included the closely related *A. sieberi* A.DC., a rare endemic species on Crete Island*. Alkanna tinctoria* is a tetraploid perennial herb mainly distributed in Greece and other Mediterranean countries (Dimopoulos *et al*., 2013, Strid, 2016). However, in countries such as Slovakia (Eliáš *et al*., 2015), Bulgaria (Petrova and Vladimirov, 2008), and Hungary (Király, 2007), *A. tinctoria* is considered as very rare. The species is of great interest to biochemists and pharmacists as its roots biosynthesize specialized secondary metabolites known as alkannin or its enantiomer shikonin (A/S) (Papageorgiou *et al*., 1999). The red extracts from roots have a long history of curative usage, dating back to the period of the Greek physician Hippocrates (Papageorgiou *et al*., 1999; Assimopoulou *et al*., 2011). Several modern investigations have shown the effectiveness of A/S in wound healing, and as antibacterial and anticarcinogenic agents (Malik *et al*., 2016; Zhang *et al*., 2019; Yan *et al*., 2019). Although the bioactivity of A/S from *A. tinctoria* and related species has been much explored, its natural genetic diversity remains poorly investigated (Ahmad *et al*., 2019). We employed ploidy-aware genotyping with RAD-seq, supported with cytogenetic investigations, to answer the following questions: *i*) Are there intra- and interspecific ploidy differences? Do the genotypic data suggest auto- or allotetraploid origin for *A. tinctoria*? *ii*) How much genetic diversity is present in *A. tinctoria* in Greece and how is this diversity distributed? *iii*) What are the factors shaping the observed genetic structure? Given the medical relevance and historical importance of *A. tinctoria* our population genomic and cytogenetic appraisal serves as a valuable foundation for understanding its genetic and phenotypic variability.

## 2. MATERIALS AND METHODS

### 2.1 Study species

Herein, we focus on two medicinally important species within the tribe Lithospermeae (Boraginaceae): *A. tinctoria* with a wide range in Greece and *A. sieberi,* as the potential outgroup species, with a very restricted range (local endemic to Crete). On Crete, the two species have partly overlapping distributions (Strid, 2016). Both species commonly occur at low elevation (<1000 m above sea level) and occupy similar habitats such as rocky, sandy, disturbed, and other open places in phrygana, xeric grasslands and coastal habitats (Strid, 2016).

### 2.2 Sampling and DNA extraction

Leaf samples of up to 11 individuals per sampling locality (Supplementary Data Table S1) were collected from 14 localities of *A. tinctoria* across three mainland regions of Greece (North, Central and South), as well as from additionally two *A. sieberi* localities found on the island of Crete (AT27, AT28) (Fig. 1). These areas roughly reflect four distinct phytogeographical regions of Greece i.e., North-East - NE, East-Central - EC, Sterea Ellas - StE, and Kriti and Karpathos – KK (Dimopoulos *et al*., 2016; Strid, 2016). The samples were either silica- or freeze-dried. Voucher specimens and /or living material (seeds) are maintained *ex-situ* at the Institute of Plant Breeding and Genetic Resources, Hellenic Agricultural Organization–Demeter (HAO-Demeter), Thessaloniki, Greece with specific IPEN (International Plant Exchange Network) accession numbers (Supplementary Data Table S1).

**Figure 1.**
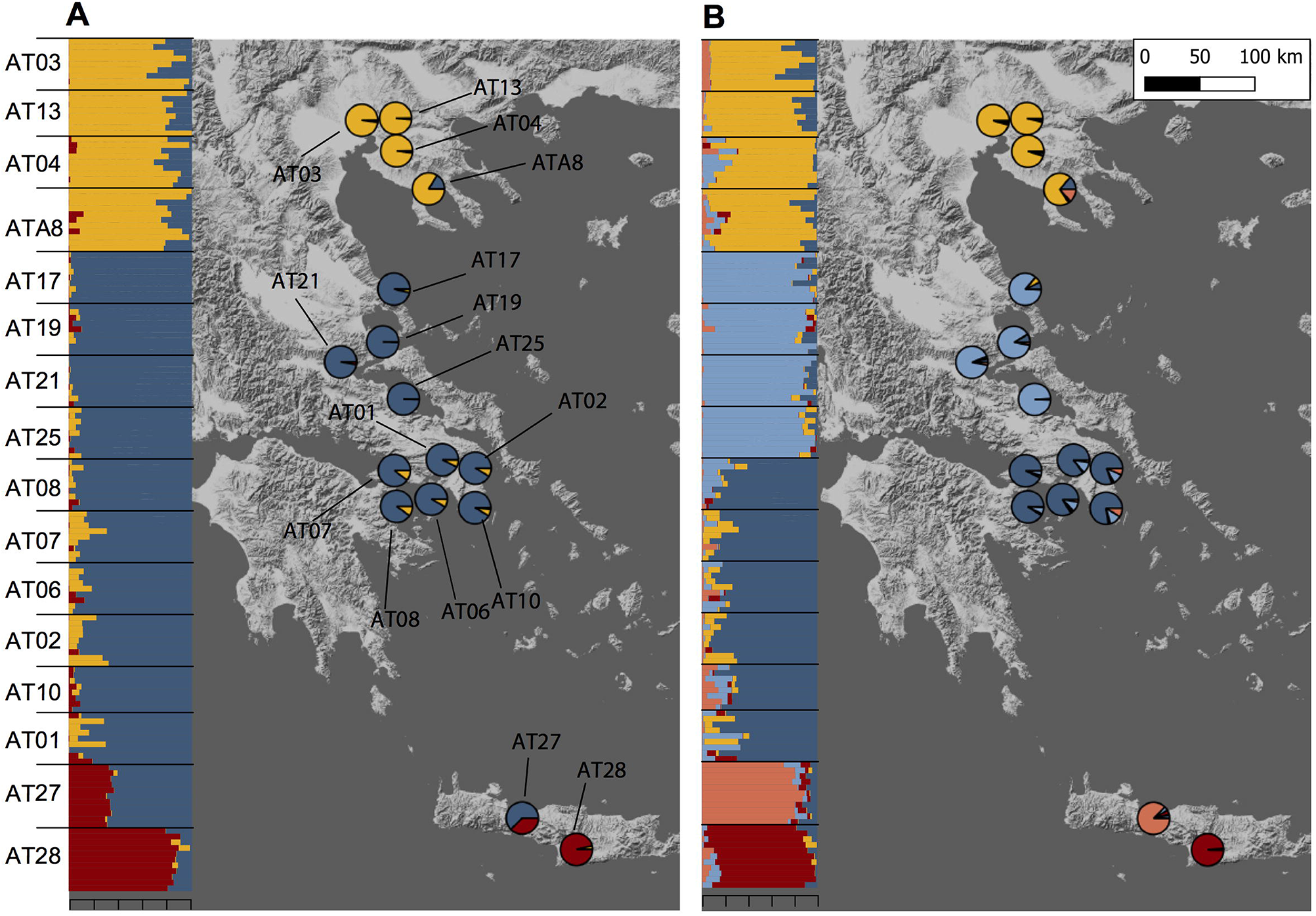
Sampling localities of *Alkanna* and its genetic structure in Greece. *Alkanna tinctoria* was sampled across Greece, and *A. sieberi* on Crete island (localities AT27 and AT28). Geographical coordinates of sampling locations are provided in Supplementary Data Table S1. Ancestry proportions inferred with TESS3 averaged for sampling locality are plotted as pie charts. Inset shows ancestry proportions from STRUCTURE as vertical bars where each vertical bar represents an individual and each color represents a genetic cluster. (A) *K* = 3, and (B) *K* = 5.

Total DNA was extracted from 40-50 mg of powdered leaf tissue using the method described in Lefort and Douglas (1999). DNA extracts were further cleaned and purified using the DNeasy PowerClean CleanUp Kit (Qiagen, Hilden, Germany) following manufacturers’ instructions.

### 2.3 Cytogenetic analysis

Fresh tissue of *A. tinctoria* was collected from three geographic regions of Greece (North, South and Central Greece), while entire plants of *A. sieberi* individuals were collected from two sampling localities (Supplementary Data Table S1) and were grown under greenhouse conditions at the Austrian Institute of Technology, Tulln, Austria. A total of ten individuals of *A. tinctoria*, each from different localities, were used for genome size assessment, plus five individuals of the outgroup species *A. sieberi* (Supplementary Data Table S1). Fresh leaf tissue was subjected to flow cytometry using about 25 mg of fresh leaf tissue, which was co-chopped together with internal standards (Galbraith *et al*., 1983), either *Pisum sativum* “Kleine Rheinländerin” (4.42pg/1C; (Greilhuber and Ebert, 1994)) or *Capsicum flexuosum* (7.44pg/1C; unpublished) in Ottós isolation buffer (Otto et al. 1981). The isolate was filtered through a nylon mesh (30-50 µm), treated with RNase in the 37°C water bath for 30 min, stained with a propidium iodide containing Ottós staining buffer (Otto et al. 1981) and stored in the refrigerator until measurement. This was performed on CyFlow flow cytometers (Sysmex/Partec, Muenster, Germany) equipped with lasers (532nm; 30mW Partec, Muenster, Germany, or 100mW Cobolt Samba, Cobolt AB, Solna, Sweden). The 1C-values were calculated based on a linear relationship between the G1/0 mean peak positions on the fluorescence axis of the standard and the objects.

Chromosome numbers were analyzed for two individuals of *A. tinctoria* originating from Northern Greece. Actively growing root meristems were harvested from young plants, pretreated with 0.002M 8-hydoxyquinoline in darkness for 2 h at room temperature and 2 h at 4°C, and fixed in 3 : 1 ethanol : acetic acid. Briefly, root meristems were hydrolyzed in 5N HCl for 20 min, washed with water, and stained with Schiff’s reagent (Merck, Darmstadt, Germany) for 1 h at room temperature in darkness. Root meristems were then squashed in a drop of 60% acetic acid under a cover slip and the preparations were analyzed using AxioPlan light microscope (Carol Zeiss, Vienna, Austria) equipped with a CCD camera. Images were captured using ZEN software (Carl Zeiss, Vienna, Austria).

### 2.4 Library preparation and RAD-seq

Preparation of the RAD-seq libraries for a total of 148 accessions (n = 8-11 per sampling locality, average 9.3; Table 2; Supplementary Data Table S1) was carried out following the protocol described in Paun et al. (2016) with modifications. Per accession 150 ng of DNA was digested with the restriction enzyme *PstI*, except for repeats where between 75 and 100 ng of starting DNA was used. After ligation of P1 adapters to digested samples overnight at 16°C, samples with different P1 barcodes were pooled together and sheared via sonication with a Bioruptor Pico (Diagenode) using three cycles of 45 s ON and 60 s OFF at 4°C. Illumina HiSeq sequencing as single-end 100 bp reads has been performed at the NGS Unit of Vienna Biocenter Sequencing Facility (VCBF; http://vbcf.ac.at/ngs/), Vienna, Austria.

**Table 1.**
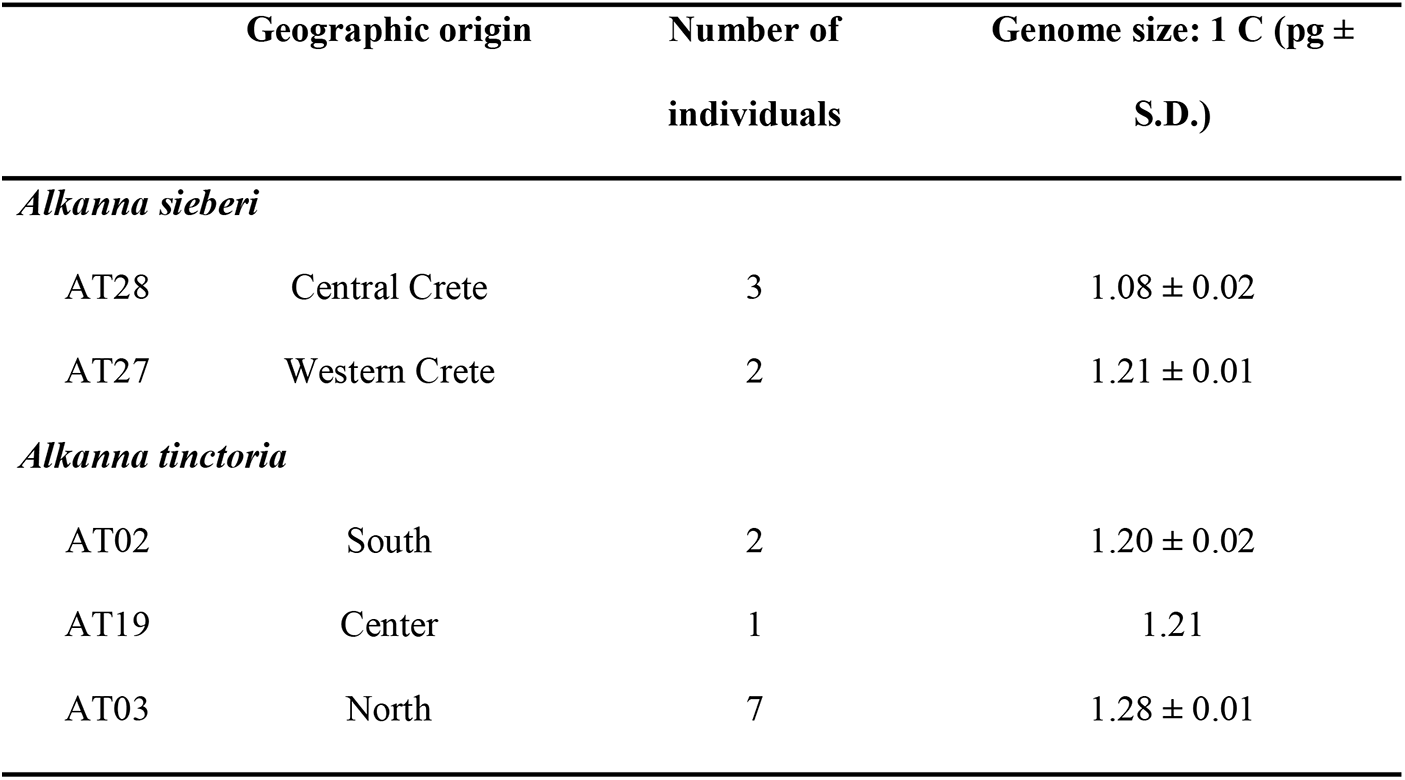
Genome size (average picogram (pg) **±** standard deviation (S.D.) of Greek *Alkanna tinctoria* and *A. sieberi* as measured by flow cytometry.

**Table 2.**
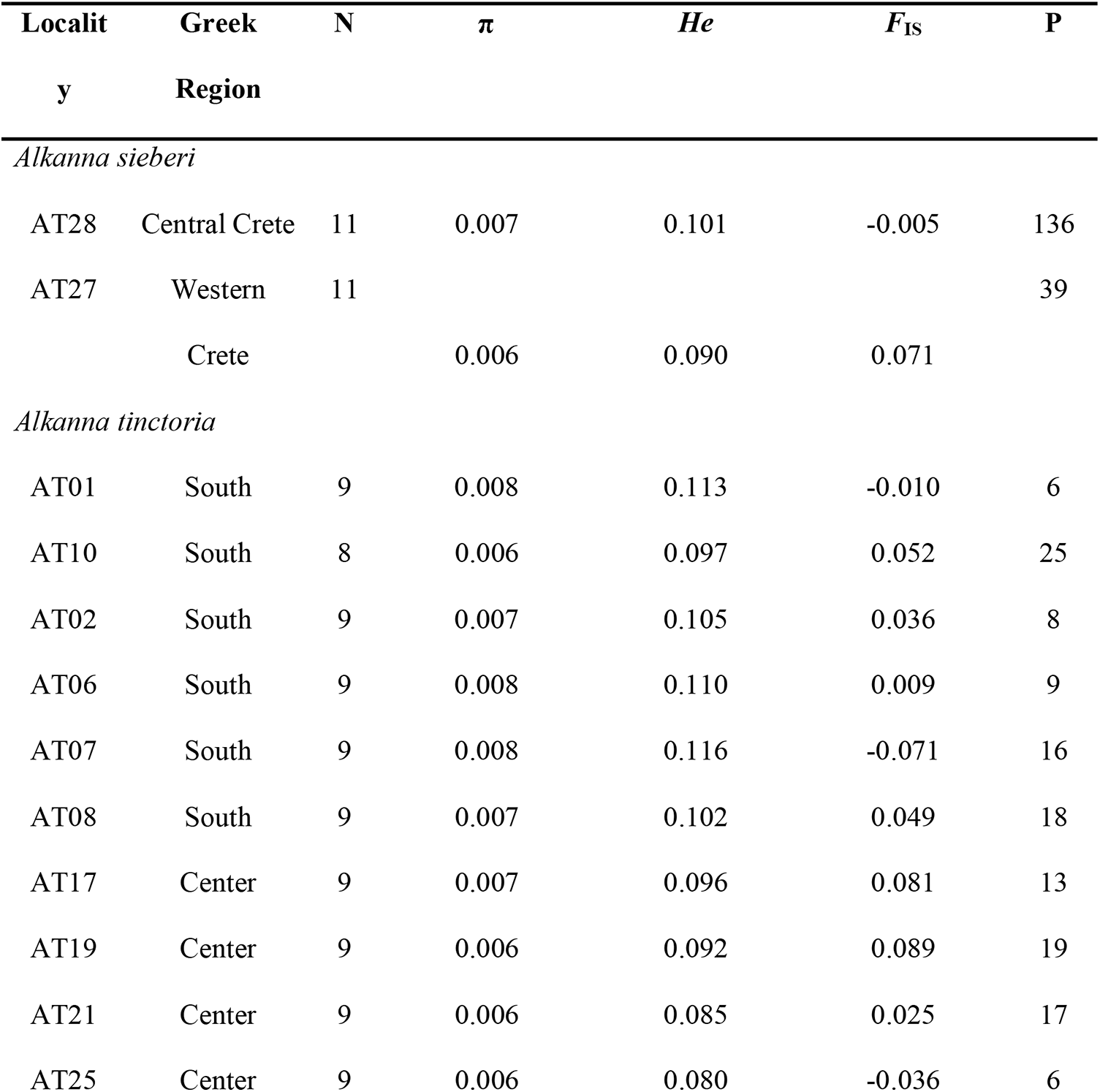
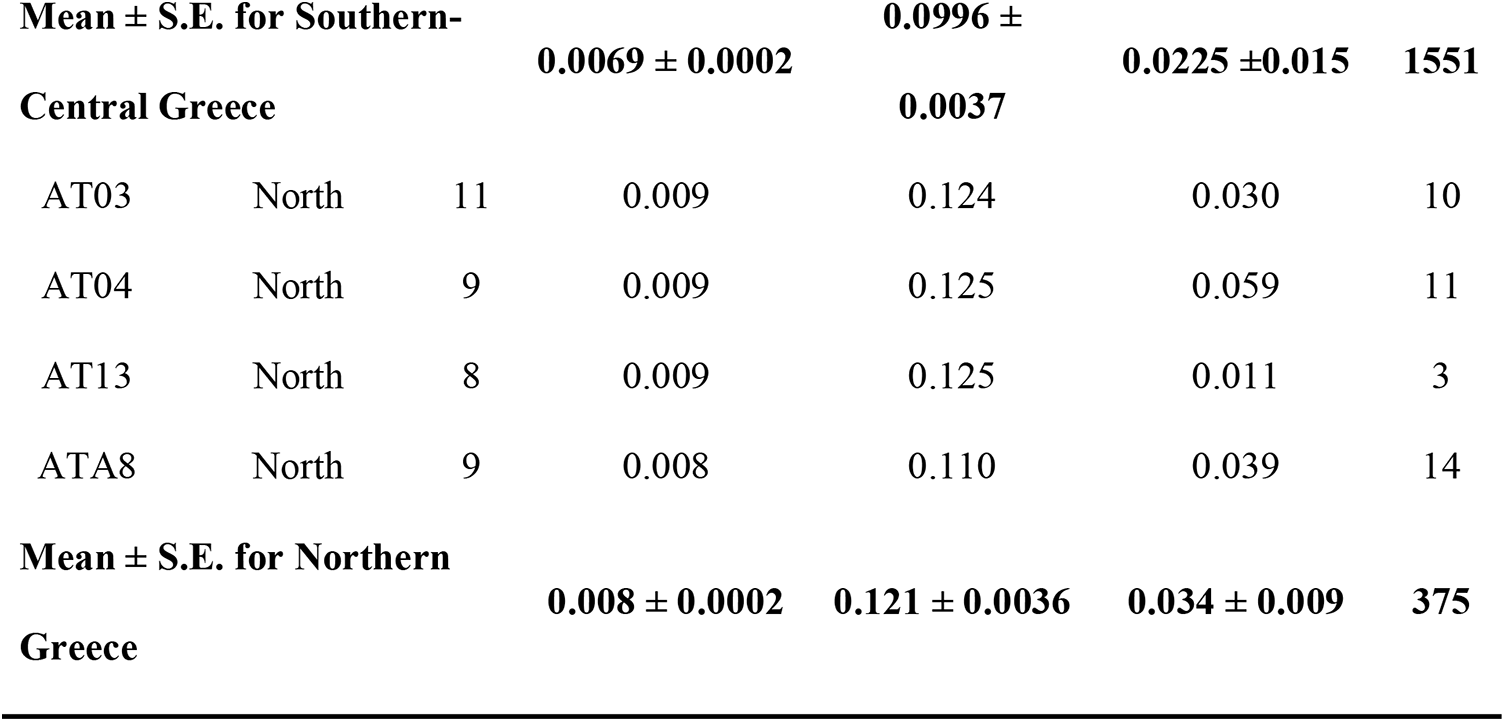
Genomic diversity in *Alkanna* populations. Summary statistics per sampling locality and mean of genetic data of two *Alkanna* species studied here. Estimates are inferred from 16,107 SNPs derived from RAD-seq data. N, Number of individuals analyzed; π Nucleotide diversity; *He*, Gene diversity; *F*_IS_, Inbreeding coefficient; P, private alleles. The statistics of genetic groups identified in *Alkanna tinctoria* by clustering-based approaches (North vs South) are highlighted in bold and represented as mean ± standard error (SE).

| Localit<br>y | Greek<br>Region | N | $\pi$ | *He* | $F_{IS}$ | P |
| --- | --- | --- | --- | --- | --- | --- |
| *Alkanna sieberi* |  |  |  |  |  |  |
| AT28 | Central Crete | 11 | 0.007 | 0.101 | -0.005 | 136 |
| AT27 | Western<br>Crete | 11 | 0.006 | 0.090 | 0.071 | 39 |
| *Alkanna tinctoria* |  |  |  |  |  |  |
| AT01 | South | 9 | 0.008 | 0.113 | -0.010 | 6 |
| AT10 | South | 8 | 0.006 | 0.097 | 0.052 | 25 |
| AT02 | South | 9 | 0.007 | 0.105 | 0.036 | 8 |
| AT06 | South | 9 | 0.008 | 0.110 | 0.009 | 9 |
| AT07 | South | 9 | 0.008 | 0.116 | -0.071 | 16 |
| AT08 | South | 9 | 0.007 | 0.102 | 0.049 | 18 |
| AT17 | Center | 9 | 0.007 | 0.096 | 0.081 | 13 |
| AT19 | Center | 9 | 0.006 | 0.092 | 0.089 | 19 |
| AT21 | Center | 9 | 0.006 | 0.085 | 0.025 | 17 |
| AT25 | Center | 9 | 0.006 | 0.080 | -0.036 | 6 |

| Mean ± S.E. for Southern-Central Greece |  |  | 0.0069 ± 0.0002 | 0.0996 ±<br>0.0037 | 0.0225 ±0.015 | 1551 |
| --- | --- | --- | --- | --- | --- | --- |
| AT03 | North | 11 | 0.009 | 0.124 | 0.030 | 10 |
| AT04 | North | 9 | 0.009 | 0.125 | 0.059 | 11 |
| AT13 | North | 8 | 0.009 | 0.125 | 0.011 | 3 |
| ATA8 | North | 9 | 0.008 | 0.110 | 0.039 | 14 |
| Mean ± S.E. for Northern<br>Greece |  |  | 0.008 ± 0.0002 | 0.121 ± 0.0036 | 0.034 ± 0.009 | 375 |

### 2.5 *De novo* assembly of loci, filtering, and polyploid genotype calling

Raw reads were first demultiplexed into sub-pools based on index reads using BamIndexDecoder (v.1.19) from Illumina2bam (available at https://gq1.github.io/illumina2bam/) with default options. Further demultiplexing into individual samples based on inline barcodes used *process_radtags* (Stacks v.1.47; Catchen *et al*., 2013), rescuing barcodes and restriction sites (*-r*) with a maximum of one mismatch, and discarding reads with low-quality scores (*-q*) and any uncalled bases (*-c*). As no reference genome is available to date for *Alkanna*, we first assembled the RAD loci with Stacks. To optimize the *de novo* assembly, we followed Heckenhauer *et al*. (2019) and varied the value of minimum reads required to create a stack (m), the maximum mismatches allows between stacks within individuals (M), and the number of mismatches allowed between sample loci (n) to maximize the number of obtained polymorphic loci including a maximum of ten SNPs and that were recovered for at least 80% of individuals. The assembly parameters were optimized on a subset of 60 individuals. The final assembly parameters chosen based on the criteria described above to produce the final catalogue were *m* = 3, *M* = 1, and *n* = 4. The RAD-seq loci from the catalog have been extracted with the *EXPORT_SQL.pl* module of Stacks. They have been further used as individual contigs to generate a synthetic reference in FASTA format.

Following the pipeline in Brandrud *et al*. (2020), the raw reads were mapped back to this synthetic reference using bowtie2 v.2.3.4.1 (Langmead and Salzberg, 2012) with default settings. Further processing (conversion to BAM, sorting by reference coordinates, and addition of read group information) was done using Picard v.2.20.6 (http://broadinstitute.github.io/picard/), and realignment around indels was performed using the Genome Analysis Toolkit (GATK v.3.81; McKenna *et al*., 2010).

We further inferred tetraploid genotypes with the GATK-based approach implemented in EBG v.0.3.2-alpha (Empirical Bayes Genotyping in Polyploids, Blischak *et al*. 2018). This pipeline is able to refine polyploid genotypes starting from flat priors and has been shown to result in negligible levels of genotype errors for both auto- and allopolyploids for intermediate to high coverages. Briefly, as recommended we employed the GATK UnifiedGenotyper for variant calling and genotype estimations with ploidy set to four. Further, we filtered the GATK generated tetraploid VCF file using *filter-VCF.R* (Blischak *et al*., 2018) to retain only biallelic SNPs present in at least 90% of individuals with a minimum quality threshold of 100 and a minimum read depth of five. The resulting polymorphic position were then used to extract base quality scores from the original BAM files using the *mpileup* command of SAMtools v.1.6 (Li *et al*., 2009; Li, 2011), to extract read counts and to estimate per site error rates with *read-counts-from-VCF.py* and, respectively, *per-locus-err.py* scripts provided with EBG (Blischak *et al*. 2018). Finally, we inferred final genotypes as alternative allele counts using the GATK model implemented in EBG.

### 2.6 Genetic diversity and population structure of Greek *Alkanna*

The summary statistics for each sampling locality was assessed by estimating within-population genetic diversity (Hs, analogue to the expected heterozygosity [*He*]) and inbreeding coefficient (*G*_IS_, analogue to *F*_IS_) in GENODIVE v.3.0 (Meirmans, 2020). In addition, nucleotide diversity (π) was estimated with a custom script (**Notes S1**). Finally, we calculated the amount of private alleles for each sampling locality as well as for the genetic groups identified by clustering-based approaches. Private alleles calculations were corrected for a small sample size using the factor ((n+1)/n) following Brandrud *et al*. (2020), where n is the number of individuals in the relevant group.

To further assess genetic relationships among the sampled individuals, we first estimated a pairwise relatedness with a method of moments as implemented in PolyRelatedness v.1.8 (Huang *et al*., 2016). This was then displayed as a heatmap of coancestry with the *heatmap.2* function in the R package Gplots v.3.1.1 (Warnes *et al*., 2009). Next, we carried out a principal component analysis (PCA) based on the covariance of allele frequencies between individuals using GENODIVE (v.3.0; Meirmans, 2020). The first three PCs were visualized as scatter plots using the R package Ggplot2 v.3.2.1 (Wickham, 2016). Only SNPs with no missing data were used for the aforementioned analyses.

We further assigned individuals to genetic groups using two clustering-based approaches: TESS3 v.1.1.0 implemented in the R package tess3r (Caye *et al*., 2016) and STRUCTURE v.2.3.4 (Pritchard *et al*., 2000). For TESS3, we used a dataset including a single SNP per RAD-seq locus (in total 7,935 SNPs) with maximum 10% missing data, whereas 1,000 SNPs were randomly selected from this dataset for STRUCTURE analyses. TESS3 is a spatially explicit clustering program and therefore it is more robust when geographic discontinuities are present in the sampling. Furthermore, TESS3 can accommodate data with thousands of SNPs and remains faster as compared to STRUCTURE. Importantly, like STRUCTURE, TESS3 can analyze polyploid data, which is otherwise currently not possible in other fast algorithms such as FastStructure (Raj *et al*., 2014) or Admixture (Alexander and Lange, 2011). In TESS3, the clustering was performed from *K1* to *K10* with ten replicates of each *K*. The final choice of *K* was based on cross validation criteria. Our objective using STRUCTURE was to confirm the individual assignment inferred by TESS3. We ran STRUCTURE with 20 replicates for each *K* value ranging from *K=1* to *K=6*, with a burnin period of 50,000, and 100,000 steps. The optimal number of ancestral populations from STRUCTURE results were determined using Evanno’s ΔK method (Evanno *et al*., 2005) as implemented in STRUCTURE HARVESTER (Earl and vonHoldt, 2012) and the replicate runs were summarized using CLUMPAK online server (http://clumpak.tau.ac.il/; Kopelman *et al*., 2015).

To quantify the strength of genetic differentiation among all sampling localities as well as among the *K* groups identified as above, we estimated pairwise *F*_ST_ with GENODIVE (v.3.0). For *F*_ST_ estimation between *K* groups, we only included those individuals where the individual assignment to respective groups was greater than 0.8 at the optimal value of *K*.

### 2.7 Spatial versus environmental contribution to genomic divergence

To disentangle the roles of dispersal limitation versus environmental drivers in shaping the genomic differentiation of *A. tinctoria*, we first identified the most important climatic variables for our target species using a gradient forest (GF) analysis in the R package gradientForest v.0.1-17 (Ellis *et al*., 2012). As a nonparametric, machine learning regression tree approach, GF evaluates the associations between genetic variants and other variables by partitioning allele frequency data at split values along the covariate gradients and defining the explained variability as split importance values. The overall importance of each covariate can then be assessed from GF plots which show the mean importance of the covariate weighted by R^2^ (Bay *et al*., 2018; Jia *et al*., 2020; Zhang *et al*., 2020). For GF analysis, we retrieved 19 bioclimatic variables for our sampling locations from the WorldClim database (Hijmans *et al*., 2005) at a spatial resolution of 2.5 arc-minutes. To avoid collinearity among covariates across our data points, the Pearson correlation coefficient between the covariates was estimated using the *cor* function implemented in the R package stats (R Core Team, 2020). We retained only one variable from all pairs having absolute correlation values greater than 0.9 using the *findcorrelation* command implemented in the R package caret v 6.0-86 (Kuhn *et al*., 2016). Finally, a total of seven covariates were used in the GF analysis. The three most important covariates identified by GF were then used in downstream analysis.

Next, we performed simple and partial Mantel analyses to test whether geographical or environmental processes contributed to the observed genome-wide divergence among *A. tinctoria* populations. The analyses were performed on pairwise genetic distances among populations as linearized *F_ST_* expressed as (*F_ST_*/*1-F_ST_*) and log-transformed geographic or environmental distances. Partial Mantel test was performed by controlling the respective geographical or environmental distance matrices. These analyses were carried out using GENODIVE with 999 permutations.

To further quantify the contribution of IBD and IBE, we employed a multiple matrix regression with randomization (MMRR) (Wang, 2013) supplemented with a commonality analysis (CA) as implemented in Cuevas *et al*. (2020). MMRR is a multiple regression method where distance matrices are used as input to assess the contribution of multiple predictor variables (*i.e.,* geography and environment) to patterns of genetic divergence. The fit of the model is assessed by R^2^ (coefficient of determination), where the statistical significance of the effect of the predictor variable (βn) and R^2^ are estimated by randomized permutations of columns and rows of the dependent variable distance matrix (*i.e*., genetic distances). However, this approach cannot take into account the multi-collinearity among predictors which can affect the estimation of regression coefficients (βn), R^2^ and p-values (Prunier *et al*., 2015). To overcome this limitation, we followed the implementation of CA suggested by Prunier *et al*. (2015) and Cuevas *et al*. (2020). This analysis decomposes the model coefficients into unique and common variance components. The unique effect represents the sole contribution of the predictor; while the common component quantifies the proportion of variances contributed by the collinearity among two or more predictor variables. Here, CA allows estimating the unique and common effect of geographical and environmental distances on genomic divergence (*F*_ST_) while accounting for collinearity among explanatory variables. We ran MMRR using pairwise *F*_ST_ as a response variable and geographical and environmental distances as predictor variables (model = *F*_ST_ ∼ GEO + ENV) with 1,000 permutations. Variance partitioning analysis was then performed using CA to quantify the common and unique effect of GEO and ENV using a R-script available at https://datadryad.org/stash/dataset/doi:10.5061/dryad.q573n5th7 (Cuevas *et al*., 2020)

In addition, we identified loci associated with environmental variables using BayPass v.2.2 (Gautier, 2015). In a nutshell, BayPass can identify loci under selection using an FST-like statistic (XtX) that explicitly accounts for the confounding effect of population structure. To do so, BayPass first estimate the covariance matrix of population-level allele frequencies (Ω matrix in Gautier, 2015), which can capture even complex demographic histories. Then BayPass detects outlier loci among variants exhibiting the highest XtX values. BayPass can also identify variants associated with population-specific covariates, including climate or phenotypic data (Gautier *et al*., 2018; Leroy *et al*., 2020). The program estimates Bayesian factors (BF) by comparing models with and without covariates. Neutral calibrations of the XtX and BF metrics were performed using pseudo-observed datasets (PODS), following the best practices for BayPass (Gautier, 2015). PODS were generated for 100,000 SNPs using *BayPass_utils.R* following the recommendations of the BayPass manual. We then used conservative (XtX > 99% quantiles and BF > 99.9%) and very conservative quantiles for this neutral simulation (XtX > 99.9% quantiles and BF > 99.9%) as thresholds to define outliers in the real dataset. All analyses were run using the standard covariate model (STD) and default parameters, and with scaled covariates, as suggested by Gautier (2015).

## 3. RESULTS

### 3.1 Genome size and chromosome counts

To rule out the possibility that the target species have different ploidy, we first estimated the genome size across multiple accessions. The genome size of *A. tinctoria* was found to vary from 1.20 to 1.28 pg (1C-value), whereas greater variation was observed among the *A. sieberi* samples, with sizes from 1.08 to 1.20 pg (Table 1). Neither species showed apparent classes of 1C values suggestive of multiple ploidies, nor were there substantial differences in 1C values of geographic relevance. This suggests that both species share the same ploidy. These results are the first reported for *A. tinctoria* and *A. sieberi*. Within the genus, the only yet published specieś genome size is 11.10 pg/1C in *A. leiocarpa* (Bou Dagher-Kharrat et al., 2013), which is the approximately 10-fold compared to our findings in *A. sieberi* and *A. tinctoria*. Finally, chromosome counts for two *A. tinctoria* samples were performed and were consistent with 2*n* = 30 chromosomes in this species (Supplementary Data Fig. S1).

### 3.2 Polyploid genotype calling supports tetratomic inheritance

After demultiplexing, RAD-seq data included on average 1.2 (SD ± 0.5) million high quality reads per individual. The optimized *de novo* assembly resulted in a catalogue of 33,126 polymorphic RAD tags of a length of 94 bp. On average, 44.0% (SD ± 8.5%) of the reads mapped back to these contigs with a final average coverage across individuals of 16× (SD ± 6×). Filtering of variants obtained from the EBG pipeline based on missing data and minor allele frequency yielded 16,107 high quality SNPs. The distribution of tetraploid genotypic frequencies against their allelic frequencies (estimated from EBG genotypes) in *A. tinctoria* showed the modes of the distribution of heterozygous genotypes (i.e., Aaaa, AAaa and AAAa) to be of similar heights in agreement with the hypothesis of tetratomic inheritance (Fig. 2; Arnold *et al*., 2015; Lloyd and Bomblies, 2016; see also discussion below).

**Figure 2.**
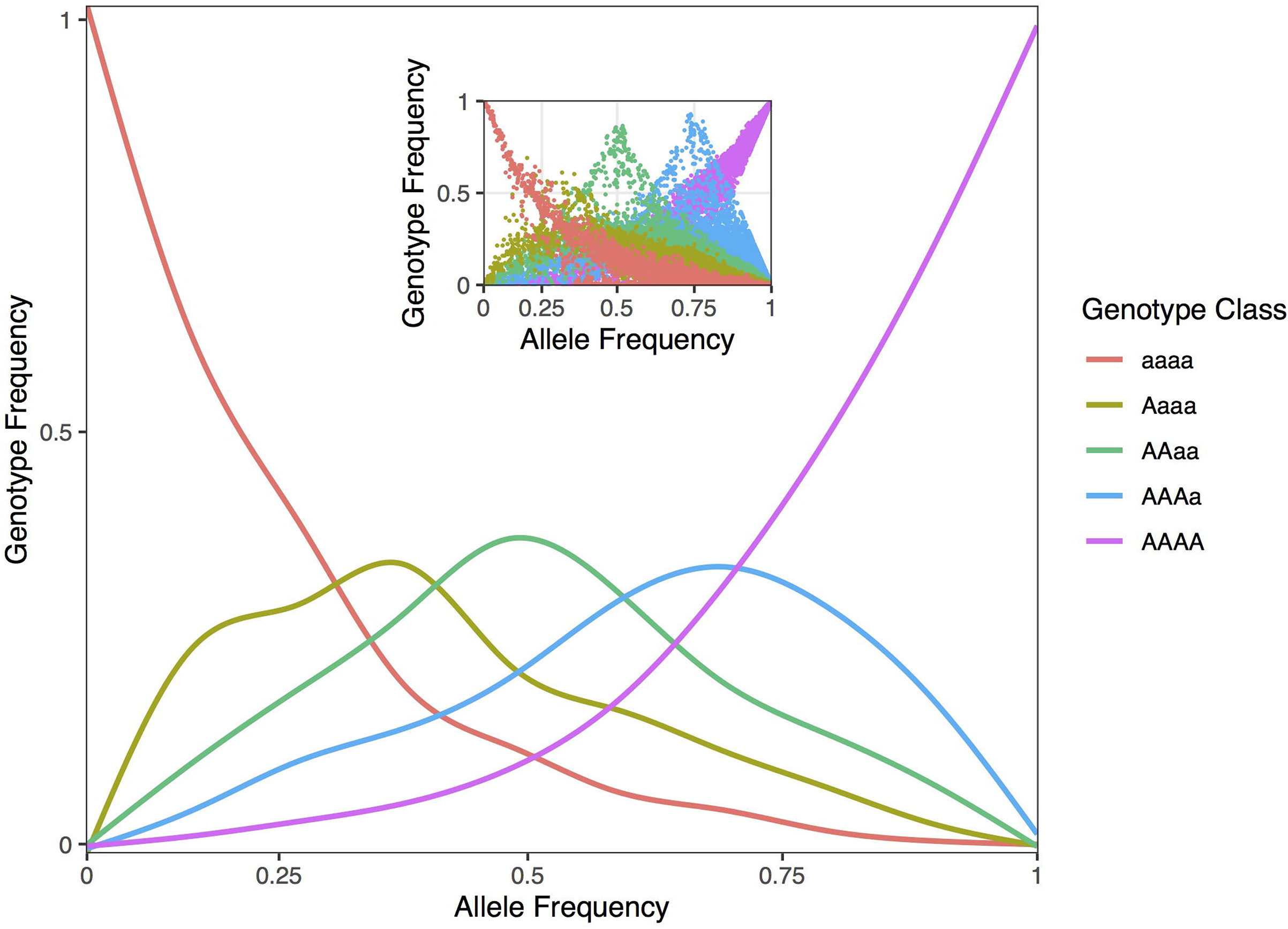
Population genomic data consistent with tetrasomic inheritance. Genotype frequencies (y-axis) as function of allele frequencies (x-axis) in *Alkanna tinctoria.* The inset shows raw plotting of genotype and allele frequencies of *A. tinctoria.* For tetrasomic inheritance, all five genotypes are expected to be observe at intermediate (0.5) allele frequency. In addition, the three middle peaks are expected to be of similar height. For disomic inheritance, all individuals are expected to have AAaa genotypes at intermediate allele frequency because of fixed differences between sub-genomes. The pattern observed here are thus, largely consistent with tetrasomic inheritance.

### 3.3 Higher diversity and lower divergence in Northern Greece

To investigate the level and distribution of genetic variation across the sampled localities, we estimated summary statistics. The genetic diversity estimates (π and *He*) revealed that the northern sampling localities harbor comparatively higher amount of diversity than the other sampled localities of *A. tinctoria* (Table 2), consistent with a larger effective population size (Ne) and more limited drift effects in the North. Nevertheless, on average diversity estimates per locality differed only slightly between the two species suggesting similar breeding systems and dispersal capacities within populations. Inbreeding coefficients (*F*_IS_) estimated for the sampling localities were all relatively close to zero (Table 2), consistent with Hardy-Weinberg expectations. Based on pairwise *F*_ST_ estimates, lower divergence levels were observed in northern localities, as compared to central and southern localities of *A. tinctoria* (Supplementary data Table S2). Highest divergence levels were observed for the locality AT25 in Central Greece, where *F*_ST_ estimates ranged from 0.144 (AT25-AT17) to 0.409 (AT25-AT28; Table 2). More surprisingly, genetic divergence between *A. sieberi* localities were also high (*F*_ST_ = 0.333); however, one of the *A. sieberi* locality (AT27) showed lower differentiation to *A. tinctoria* populations than to the second (AT28) locality of *A. sieberi* (Supplementary Data Table S2).

### 3.4 North-south population structure in Greek *Alkanna*

The relatedness-based heatmap allows to identify three main genetic groups (Fig. 3A). Two main groups were specific of *A. tinctoria* hereby called North and Southern-Central. Within the latter group, the analysis allows to find evidence of substructure, both at the regional scale (Central vs. Southern) and even sometimes, more locally, between the different sampling sites (*e.g.* AT25, Fig. 3A). The third main group consisted of the *A. sieberi* locality from central Crete (AT28), which showed the least relatedness to all *A. tinctoria* accessions (Fig. 3A). However, interestingly, all individuals from the western Cretan locality (AT27) that have been initially identified as *A. sieberi* showed intermediate relatedness to the individuals of *A. tinctoria* from Southern-Central group, and to *A. sieberi* from the central Cretan locality (AT28), therefore suggesting interspecific admixture. One accession from *A. tinctoria* from South Greece (AT01) shows also increased co-ancestry with *A. sieberi* (indicated with an arrow in Fig. 3A), suggesting recent gene flow between *A. tinctoria* and *A. sieberi*.

**Figure 3.**
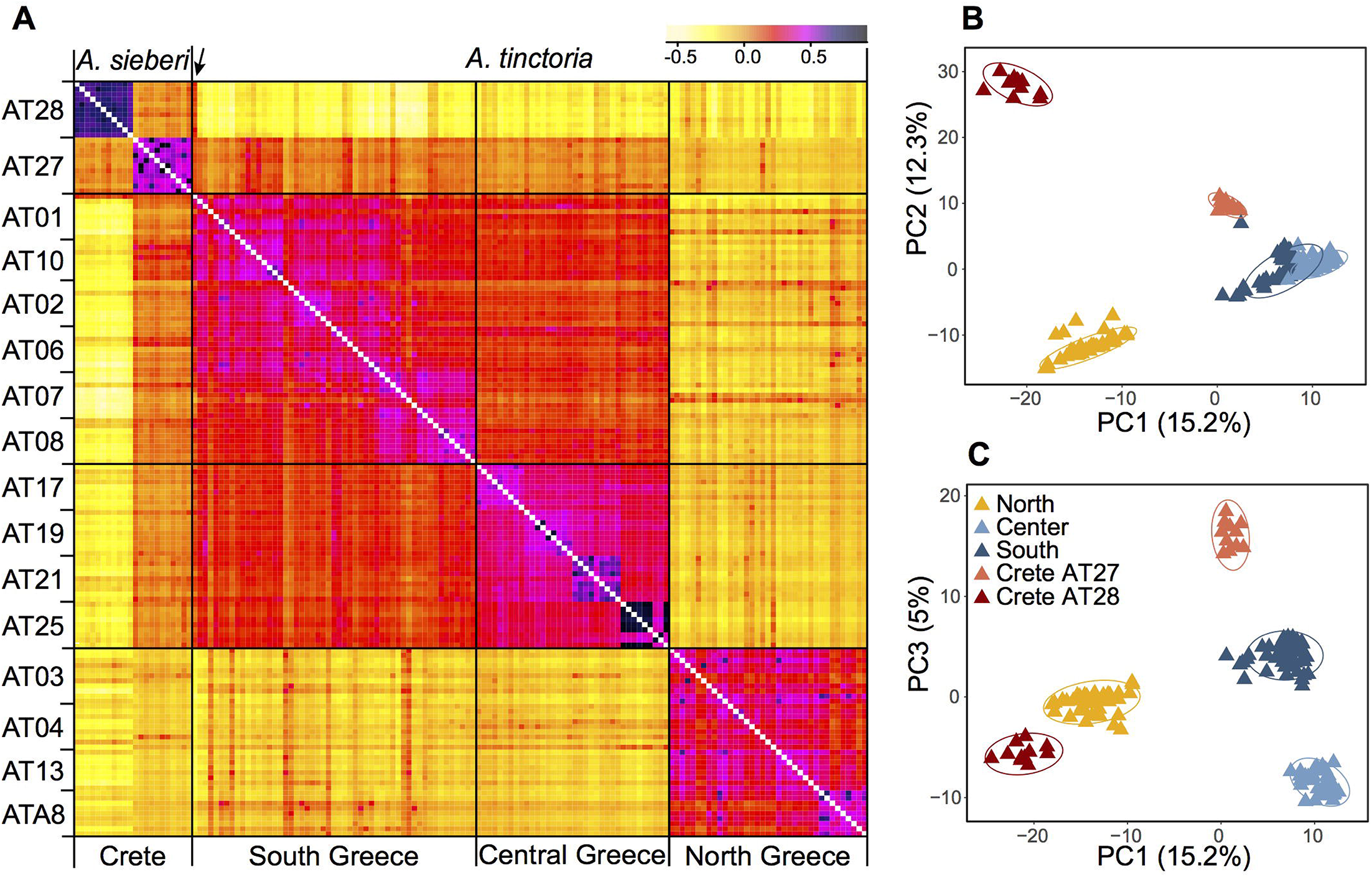
Genomic variation in *Alkanna*. **A.** Heatmap of pairwise relatedness between 148 individuals from 16 different localities across Greece based on 16,107 SNPs. Regional origins of individuals are given below the heatmap and each population is indicated on the left side of the heatmap. Darker colors represent higher relatedness, according to the legend included on the top right side. For improved color resolution, the estimates on the diagonal were excluded. An arrow points to a potentially admixt individual in population AT01, that is discussed in the text. **B-C.** Genetic structure in the Greek *Alkanna* populations inferred based on EBG-derived tetraploid genotypes (n = 148) as revealed by principal component analysis (PCA).

To further investigate population structure, we performed PCA (Fig. 3B-C), which highlighted consistent patterns with the previously identified clusters in the co-ancestry heatmap. The first two PCs, which together explained 27.6% of the total variance, primarily distinguished individuals of *A. sieberi* from the central Cretan locality (AT28), and those of *A. tinctoria* from the North on one side and from the Southern-Central localities on the other. The individuals from the western Crete population (AT27), plus the accession from AT01 already identified as admixed based on the co-ancestry heatmap, were located between the *A. tinctoria* Southern-Central group and the cluster formed by *A. sieberi* accessions from AT28 (Fig. 3B), again consistent with admixture. The central Greek localities were well resolved from Southern Greece in particular by PC3, but this axis explains a lower proportion of the total variation (5%, Fig. 3C).

We then used Bayesian clustering to further investigate these patterns of genetic structure. For that purpose, we used the spatially-explicit TESS3 and the non-spatially explicit, albeit widely used, STRUCTURE programs. Both the cross-validation score as implemented in TESS3 and the use of the Evanno’s ΔK statistic supports *K* = 3 as the best *K* value (Supplementary DataFig. S2), suggesting that the dataset can be best explained by three main genetic clusters. For the optimal structure (*K* = 3), the clustering of individuals is consistent with all previously described analyses (*i.e.,* presence of three genetic groups), including the likely interspecific admixed nature of AT27 (Fig. 1A). Since heatmap and PCA showed substructure within the southern-central group, we further evaluated the individual memberships at K values from *K*= 4 to *K* = 5. As expected, at these values of *K*, a further separation of Southern and Central *A. tinctoria* was observed (Supplementary data Fig. S3), whereas eastern Crete accessions of *A. sieberi* showed signatures of admixture between the Southern-Central group and the Central Crete population at *K* = 4 (Supplementary data Fig. S3) and formed an independent group at *K* = 5 (Fig. 1B).

Finally, to evaluate the strength of genetic differentiation between the identified groups, we estimated *F*_ST_ between the three genetic pools (*K* = 3) with well assigned individuals (q > 0.80). We observed a significant genetic divergence between the three groups (p = 0.001). Genetic differentiation was highest between AT28 and mainland groups (North: 0.29 and Southern-Central: 0.31) and lowest between Southern-Central and North group (0.14).

### 3.5 Geographical and environmental influences on genomic divergence within *Alkanna tinctoria*

To assess the role of environmental and geographical variables in the onset of genetic differentiation, we first identified the most important climate-related covariates using a random forest-based approach called gradient forest (GF). Among the seven non-collinear covariates tested with GF, the top three covariates were related to precipitation, including precipitation of coldest quarter (BIO19), precipitation of warmest quarter (BIO18), and annual precipitation (BIO12) (Fig. 4A). We then used the top three climatic variables identified by GF in Mantel statistics and genotype-environment association analysis. The pairwise genetic distances (*F*_ST_/(1-*F*_ST_) between populations were strongly correlated with pairwise geographical distances (R^2^ = 0.49, p = 0.001). We also observed a milder, but still significant association between the genetic and the environmental distance matrices (R² = 0.24, p = 0.002) (Fig. 4B-C). In multiple comparisons where we controlled for either distances, the geographic distances were strongly and highly significantly associated with genetic distances (Mantel’s *r =* 0.63, p = 0.001), whereas the association with environmental distances was weaker albeit significant (Mantel’s *r =* 0.31, p = 0.020). The multivariate model (MMRR) including both predictors (*F_ST_* ∼ GEO + ENV) explained 55% (R^2^ = 0.55) of the genomic variation within the Greek *A. tinctoria* (Table 3). Consistent with the results from the partial Mantel analysis, in MMRR, the geographic distances showed a significant and a higher β weight (β_GEO_ = 0.037, p = 0.002, Table 3), explaining 75% of total variation in the model. A considerable proportion of this total variability was assigned as unique (Unique 54 %; Common 21%, Table 3). The environmental distances had a lower β weight (β_ENV_ = 0.025, p = 0.003), explaining 45% of the variation in the model (Unique 24%, Table 3). Taken together, these results suggest that IBD has a stronger contribution to the overall genetic differentiation among *A. tinctoria* populations as compared to IBE.

**Table 3.** Summary of multiple matrix regression with randomization (MMRR) and commonality analysis (CA). Pairwise genomic divergence (*FST*) between 14 sampling localities of *Alkanna tinctoria* as response variable and geographical (GEO) and environmental (ENV) distance matrices as explanatory variables. Unique, U; Common, C; and Total, T variance partitioning of each explanatory variable contributing to genomic divergence. The proportion in parenthesis represent the percentage contribution of each predictor to the total variance explained by the model and calculated from CA coefficient as: (Unique, Common or Total/R^2^)*100.

| **Model:** $F_{ST} \sim \text{GEO} + \text{ENV}$ | | | | $R^2 = 0.55$ |
| --- | --- | --- | --- | --- |
| Predictor | $\beta$ | Unique (U) | Common (C) | Total (T) |
| GEO | 0.037 * | 0.29 (54 %) | 0.11 (21%) | 0.41 (74%) |
| ENV | 0.025 * | 0.13 (24 %) | 0.11 (21%) | 0.25 (45%) |

\*significant at p = 0.001

**Figure 4.**
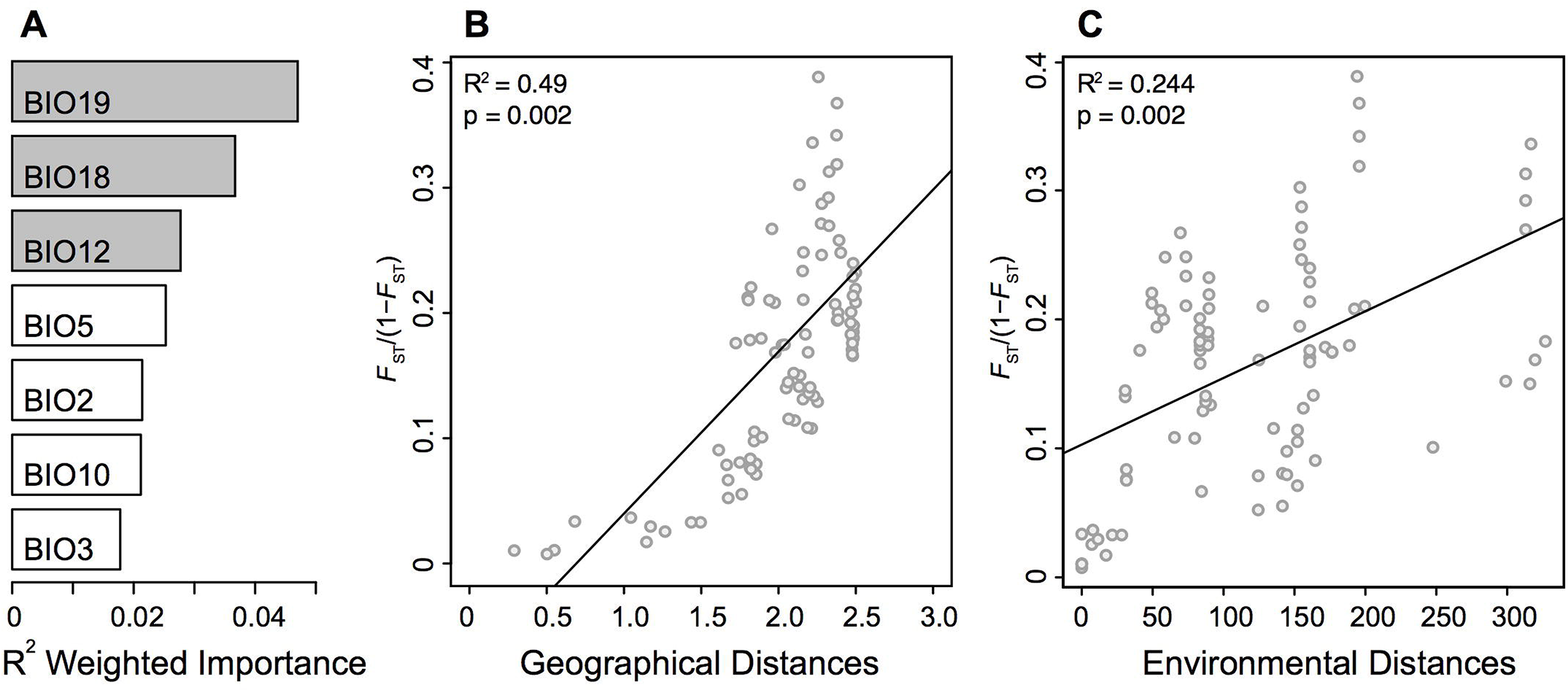
Isolation by distance (IBD) and Isolation by environment (IBE). Influence of geographical and environmental covariates on genetic differentiation in Greek populations of *Alkanna tinctoria*. **A.** R^2^ weighted importance of environmental covariates that explains genetic gradients. Top 3 environmental covariates are given in grey. Relationship of genetic distances (*F*_ST_/(*1-F*_ST_) and (**B**) log-transformed geographical distances, or (**C**) environmental distances.

### 3.6 Genotype-environment associations

Although the overall population structure appears more explained by migration-drift equilibrium, some loci could be affected by selection either directly, or through linkage. We therefore performed a *F*_ST_ outlier scan and a genotype-environment association analysis. To identify non-neutral markers, we performed a classical genome scan using the XtX statistics, an *F*_ST_ analog which explicitly accounts for the confounding effect of population structure including IBD. Using BayPass, we identified 107 SNPs (0.7% of all SNPs) with conservative criteria, including 17 (0.11%) using a more conservative criterion. Then, under BayPass, we assessed associations of three top bioclimatic variables and identified between up to 0.14% of the SNPs with BF > 99.9% depending on the variable. Interestingly, among the 107 previously detected *F*_ST_ outliers, several SNPs were associated to precipitation: six SNPs were associated with annual precipitation (BIO12; BF > 99.9%, XtX > 99%), including one SNP among the 17 strongest outliers. Similarly, six and four SNPs were associated with precipitation of warmest quarter (BIO18) and of coldest quarter (BIO19), including three and one SNPs among the 17 strongest outliers, respectively (Supplementary Data Fig. S4). Among all these outliers, three SNPs were associated with both BIO12 and BIO19 (Supplementary Data Fig. S4). Clinal variation of allele frequencies across each clinal gradient follows linear trends as shown in Supplementary Data Fig. S5.

## 4. DISCUSSION

RAD-seq is being commonly employed to study the intra- and interspecific genomic divergence, to investigate the levels of genomic diversity and to resolve phylogenomic relationships of diverged lineages (Paun *et al*., 2016; Rochette and Catchen, 2017; Brandrud *et al*., 2020). However, the use of RAD-seq in non-model polyploid species is still challenging due to difficulties of inferring allele copy numbers (Clevenger *et al*., 2015; Wagner *et al*., 2020). Thus, only recently, RAD-seq has started to be used for analyzing population structure of non-model polyploid plant species (Zohren *et al*., 2016; Brandrud *et al*., 2017). Here, using an appropriate ploidy-aware genotype calling approach (Blischak *et al*., 2018; Brandrud *et al.,* 2020), we investigated the main drivers of genomic diversity and divergence of *A. tinctoria* - a species of particular interest in the traditional and modern pharmacopoeia (Assimopoulou et al., 2011; Papageorgiou et al., 1999), jointly analyzed with two populatons of a potential closely related outgroup species, *A. sieberi*. Overall, we detect a marked regional structure along a latitudinal gradient and highlight an important role for physical distance between the investigated sites in shaping genetic diversity. However, we also identified loci with variation related to ecological pressures, in particular related to precipitation, suggesting that both levels can be at play in shaping the population structure in *A. tinctoria*.

### 4.1 Limited genome size variation and fixed dysploidy at the tetraploid level

Considering all available evidence, including the newly described chromosome number (2*n* = 30), previous microsatellite analyses where up to four alleles were observed per individual (Ahmad *et al*., 2019), and previously reported base chromosome numbers in the genus (*x* = 7 most frequently reported, but also *x* = 10, 11 and the derived secondary base number of *x_2_* = 15; Coppi *et al*., 2006; Weigend *et al*., 2016), we inferred that *A. tinctoria* is a dysploid at the tetraploid level (4*x* + 2) derived from lineages/species with *x* = 7. Indeed, *x* = 7 is one of the widespread in Lithospermae (Weigend *et al*., 2016). A tetraploid state is also strongly supported by the present genomic analyses, in particular by the similar frequencies of Aaaa, AAaa and AAAa genotypes when the data is scored as tetraploid (Fig. 2). On the other hand, the relatively consistent genome size estimates within *A. tinctoria*, and the lack of substantial genome size differences between *A. tinctoria* and *A. sieberi* (Table 1), suggests that the sampled accessions of these two species represent a single ploidy. This is in agreement with previous karyological findings for *A. tinctoria* (Grau, 1968; Kamari and Papatsou, 1973; Luque, 1990; Markova, 1995), supporting a single ploidy across its entire distribution range.

In tribe Lithospermae, several tetraploid species have been identified (Luque, 1990; Korbecka *et al*., 2003; Coppi *et al*., 2006; Kolarčik *et al*., 2014; Weigend *et al*., 2016). Moreover, both allopolyploids and autopolyploids have been described in this tribe (Luque and Valdes, 1984; Kolarčik *et al*., 2014; Kolarčik *et al*., 2015; Weigend *et al*., 2016). In the present study, two lines of genomic evidence led us to suggest that *A. tinctoria* might be an autopolyploid. First, in allotetraploids, especially those with well differentiated parents and disomic inheritance, the proportion of duplex heterozygous genotypes (*e.g*., AAaa) is expected to be significantly higher than other classes of heterozygous genotypes (*e.g*., AAAa and Aaaa). At the other end of a continuum, autotetraploids with tetrasomic inheritance are expected to show the reversed trend. In *A. tinctoria,* we observed that triplex heterozygous genotype class (i.e., AAAa and Aaaa together) is approximately two-fold more frequent than the duplex heterozygous genotypes (Fig. 2). Second, for loci with intermediate allele frequency, allopolyploids are expected to be largely represented by AAaa genotypes because of fixed differences between divergent subgenomes (Lloyd and Bomblies, 2016). In contrast, in autopolyploids, all five genotype classes are expected to be observed around this allele frequency (Lloyd and Bomblies 2016). When *A. tinctoria* genotype proportions were plotted against allele frequencies, all five classes of genotypes could be seen at intermediate allele frequency (Fig. 2). This distribution is comparable to that described for autotetraploid *Arabidopsis arenosa* (Arnold et *al.*, 2015) and it is expected under a tetrasomic mode of inheritance (see Fig. 1c and 1d in Lloyd and Bomblies, 2016). However, we note that in depth cytological investigation would be required to confirm the hypothesis put forward here.

### 4.2 Spatial structure in Greek Alkanna is associated to a latitudinal gradient

Genetic diversity within localities of *A. tinctoria* was moderate and mostly homogenous across the sampling range, with higher values in the North (Table 2). Although not directly comparable, a previous study also reported moderate albeit similar levels of genetic diversity at inter simple repeat markers (ISSR) in three other *Alkanna* species (Semerdjieva *et al*., 2020). The relatively higher genetic diversity, and lower genetic differentiation between sampling localities (Table 2 and Supplementary Data Table S2) observed in the North could be a result of higher connectivity of stands and larger *Ne* leading to lower effect of genetic drift as compared to central Greece where *A. tinctoria* has a much more fragmented distribution. During field visits, we observed an absence of *A. tinctoria* from several regions located around East-Central Greece (*e.g.*, Lárisa, Magnisia, and Fthiotida) probably due to a change of management practices (conversion into agricultural lands), or due to expanding urbanization over natural habitats. Reduced connectivity of suitable habitats could lead to genetic isolation (Toczydlowski and Waller, 2019) and thus expose populations to greater effects of genetic drift. Consistent with an increased effect of genetic drift in more isolated localities, we observed highest amount of *F*_ST_ as well as reduced genetic diversity at locality AT25 in central Greece which appeared disconnected from other populations by vast agricultural landscape.

Genetic structure analyses identified two divergent *A. tinctoria* clusters, localized around Northern and, respectively, Southern-Central Greece, with a weaker subdivision between South and Central accessions (Fig. 1 and 3), consistent with a latitudinal structuring of regional genetic variation. The strength of divergence was further confirmed by a significant *F*_ST_ (0.14, p = 0.001) estimated between these main genetic pools. Although investigating different regions and spatial scales, other studies also found evidence for significant genetic structure in other *Alkanna* species (Wolff *et al*., 1997; Semerdjieva *et al*., 2020)

Genetic structure, such as that observed in *A. tinctoria*, is possibly explained by the establishment of populations from multiple glacial refugia as have been reported for other Balkan and Mediterranean regions (for a review see Nieto Feliner, 2014). However, this might be a less likely explanation for the Greek *A. tinctoria,* as currently available phylogeographic or phylogenetic studies on plants (Lakušić *et al*., 2013; Jaros *et al*., 2018; Janković *et al*., 2019) from Greek part of Balkan do not provide consistent indications for a North-South genetic differentiation (but see Fassou *et al*., 2020). But what might have been the forces giving rise to the North-South clustering in Greek *A. tinctoria*? Although not explicitly tested in the current study, the identified genetic structure in mainland localities of *A. tinctoria* mirrors the Greek phytogeographic zones suggested in Flora Hellenica (Dimopoulos *et al*., 2013, 2016; Strid, 2016). For example, the northern lineage of *A. tinctoria* belongs to the North East (NE) phytogeographic region, whereas the majority of the southern-central localities pertain to the Sterea Ellas (StE), except two populations which were collected at the boundary between East Central (EC) and StE (AT17 and AT19), and between Peloponnisos (Pe) and StE (AT07 and AT08). Botanical exploration and biosystematics studies have shown consistent discontinuities in species composition, richness, and endemism patterns between StE and NE (Dimopoulos *et al*., 2013, 2016; Strid, 1996, 2016). These phytogeographical regions take into account biogeography and have been mainly defined by network biogeography (Kougioumoutzis *et al*., 2017), or comparative floristics (Strid, 1996). Hence, these divisions reflect the paleogeographic history of those regions and dispersal limitations of organisms owing to geographic as well as ecological barriers. In the case of Greece, this could include the complex geological history of the Aegean region (*e.g*., fragmentation of mainland and subsequent island formation including the Aegean archipelago), topographic heterogeneity (numerous mountain massifs and mountain ranges separated by rivers, valleys, plains and lowlands), varied climatic (descending rainfall and ascending temperature gradients from north to south) and soil features (Dimopoulos *et al*., 2016; Perlès, 2016). Hence, these phytogeographic regions represent a conglomerate of many factors (including geological, geographical and environmental covariates), which might have shaped the observed genetic differentiation in *A. tinctoria*.

### 4.3 Dispersal limitation as a primary source of genetic divergence in *A. tinctoria*

The high relatedness of geographically proximal individuals, as well as the above-mentioned clustering patterns (Fig. 1 and 3), suggest that IBD might have played a major role in shaping the observed genetic differentiation. Indeed, we inferred a significant relationship between geographical and genetic distances even after controlling for climatic covariates (Fig. 4B, Table 3). In *A. tinctoria*, seed dispersal is limited to very short distances (Ulrych and Szabóová, 2010), with pollen exchange likely promoting almost exclusively gene flow between localities. Although no detailed study has been carried out on pollination in *A. tinctoria*, existing evidence suggests that pollination is performed by bees (Ulrych and Szabóová, 2010; Dafni *et al*., 2012), insects with limited potential for long distance dispersal, especially in areas with complex topography, such as Greece, where plains are separated by mountains and sea. Such a pattern could therefore be consistent with isolation by distance, *i.e.* a migration-drift equilibrium hypothesis. Furthermore, the southern Balkan Peninsula has been recognized as an important glacial refugium for Mediterranean biota (Médail and Diadema, 2009), opening the possibility that *A. tinctoria* existed in this region for a long period. A long local history together with limited dispersal, patchy distribution and topographic barriers, could have led to the observed pattern of IBD.

### 4.4 Climate as a second source of population structure in *A. tinctoria*

Climatic variables have been implicated as strong selective factors in plants (Joshi *et al*., 2001). Understanding the association of climatic covariates with the observed genetic divergence can prove useful in determining a species’ ability to persist against varying climatic scenarios (Jia *et al*., 2020). Our study identified significant associations between environmental variables and genetic differentiation after controlling for spatial covariates (Fig. 4C, Table 3). Specifically, GF analysis indicated that the genetic structure across our sampling localities is strongly associated with precipitation characteristics (Fig. 4A). The Mediterranean climate is mainly characterized by hot, dry summers and cold, comparatively wet winters (Joffre *et al*., 1999). In this environment, the precipitation regime imposes strong constraints on plants and can act as an important evolutionary driver (Galmés *et al*., 2007). *Alkanna tinctoria* is habitually confined to sandy soils and rocky places, xeric grasslands and coastal habitats, highlighting the sensitivity to water availability, and autumnal wetter periods have been shown to coincide with rigorous plant growth for this species (Pluhár *et al*., 2001). Moreover, rainfall during late winter is important for early spring flowering in many plant species (Moore and Lauenroth, 2017). Therefore, precipitation characteristics appear to play an important role in promoting local adaptation and shaping genetic differentiation in *A. tinctoria.* Additional support for this hypothesis is also offered by the identification of outlier loci associated with rainfall (Supplementary Data Fig. S4). Although the identified loci are few and do not entirely deviate from the neutral expectations, some of these loci show substantial allele frequency changes associated with the respective covariates (Supplementary Data Fig. S5), pointing to adaptive genetic differentiation.

### 4.5 Genomic divergence and introgression in mainland-island congeners of *Alkanna*

Our investigation of genetic differentiation detected a high genetic differentiation between mainland *A. tinctoria* and *A. sieberi* accessions from one of the two sampled Cretan localities (AT28; Fig. 1 and 3; average *F_ST_* = 0.271, p = 0.001) after excluding putatively admixed individuals. Phylogenetic and historical biogeographical studies have shown that the diversification in islands mainly occurred via vicariant speciation, that is taxa become isolated owing to formation of barriers (Comes *et al*., 2008; Cellinese *et al*., 2009; Poulakakis *et al*., 2015; Sfenthourakis and Triantis, 2017; Jaros *et al*., 2018). However, secondary colonization events and gene flow between divergent populations from mainland or neighboring islands during sea level fluctuations and recurrent land bridge formation have also been proposed in other species (Li *et al*., 2010; Zhai *et al*., 2012). Due to lack of dated phylogenomic studies in genus *Alkanna* and the absence of samples of *A. tinctoria* from Crete do not allow us to predict whether the observed genetic split between mainland-island congener is a consequence of strictly vicariant divergence, or resulted from the colonization event(s), as the later have been hypothesized for some Cretan (sub)species (Cellinese *et al*., 2009; Jaros, Tribsch, & Comes, 2018).. In future, a more wide sampling of each taxon is needed to understand whether the observed pattern follows the allopatric model of vicariant divergence, or other processes (*i.e.,* post-fragment colonization) are also at play.

Intriguingly, in addition to genetic divergence between mainland-island congeners, our data identifies significant introgression between the divergent southern lineage of *A. tinctoria* and the Central population from Crete. We identified via traditional and Bayesian clustering that the investigated accessions from AT27 showed an admixed ancestry between the two species (Fig. 1 and 3). In addition, AT27 formed its own independent clade on the third PC, as well as in the Structure and TESS3 results at K = 5 (Fig. 1B), but showed comparatively lower heterozygosity. In turn, this implies that the admixture event might be old and further point towards a complex demographic history where the admixed population might have experienced bottlenecks during colonization phase resulting in the reduction of genetic diversity.

### 4.6 CONCLUSION

In the present study, we provide evidence that only dysploid tetraploid *A. tinctoria* and *A. sieberi* are prevalent in Greece, the main distribution area for these species. Our further population genetic inferences start from ploidy-aware genotype calling and successfully employ a strategy that can be extended to other polyploid non-model plant species, for example in order to develop hypotheses regarding mode of inheritance of polyploids. Overall, we found a moderate intrapopulation genetic diversity in *A. tinctoria*, but significant regional differentiation, together with clear evidence of admixture between mainland and island congeners. Furthermore, the strong genetic structure uncovered is in line with native phytogeographical divisions, and appears mainly shaped as a result of dispersal limitations and drift, but also of isolation by environment, with precipitation as the main contributing factor. Finally, the genomic variation revealed here can be used to further inform and predict the structure of phenotypic variation (*e.g*., alkannin and shikonin production) in this important medicinal plant, prompting to future studies.

## SUPPLEMENTARY DATA

Supplementary data consist of the following. Figure S1: Chromosomes of *Alkanna tinctoria.* Figure S2 A-B: Cross validation score (A) and ΔK (B) showing optimal K identified by TESS3 and STRUCTURE HARVESTER. Figure S3: Genetic structure in studied *Alkanna* populations inferred based on EBG-derived tetraploid genotypes. Figure S4 A-C: Outlier analysis of genotype-environment association (BayPass) of *Alkanna tinctoria* (A) BIO12 (B) BIO18 (C) BIO19. Figure S5 A-C: Changes in allele frequencies of loci identified by BayPass along the environmental gradient (A) BIO12 (B) BIO18 (C) BIO19. Table S1: Geographic coordinates of localities included in this study. Table S2: Pairwise *F_ST_* among sampling localities. Notes S1: Custom function used to calculate pi from EBG output file

## FUNDING

The research was funded by the European Union’s Horizon 2020 Research and Innovation Program (MicroMetabolite, grant agreement no. 721635).

## ACKNOWLEDGEMENTS

This article is dedicated to Prof. Christian Lexer, a wonderful scientist and mentor that passed away too early. We thank Angela Sessitsch and our partners in MICROMETABOLITE project for obtaining the funding, Eleni Maloupa, Andreana Assimopoulou and Nikolas Fokialakis for provision of plant material used in this study, Karin Hansel□Hohl for her support in the wet lab, Antonielli Livio and Silvia Madritsch for initial help with Bash and R scripts, and Paul Blischak for advices on EBG implementation.

## DATA ACCESSIBILITY

Raw Illumina reads are available from NCBI’s Short Reads Archive under the BioProject number PRJNA705074.

## AUTHOR CONTRIBUTIONS

MA, EMS, CL and OP planned and designed the research, NK collected samples, ET and HS performed flow cytometry and chromosomal counts. MA prepared RAD-seq libraries and analyzed the data with support from OP and TL. MA, EMS, TL and OP interpreted the results, and MA and OP wrote the manuscript with the input from all authors.

## Notes

### Competing Interest Statement

The authors have declared no competing interest.

