## Supplementary data for "Spatial and ecological drivers of population structure in *Alkanna tinctoria* (Boraginaceae), a polyploid medicinal herb"

**Table of Contents:**

| **Table S1** | Page 2 |
| --- | --- |
| **Table S2** | Page 3 |
| **Figure S1** | Page 4 |
| **Figure S2** | Page 5 |
| **Figure S3** | Page 6 |
| **Figure S4** | Page 7 |
| **Figure S5** | Page 8 |
| **Notes S1** | Page 9 |

| **Table S1.** Acronym, number of individuals included in the RAD-seq analyses (N), voucher and/or living material information with IPEN (International Plant Exchange Network) accession number, and, and geographic coordinates of localities included in this study. | | | | | |
| --- | --- | --- | --- | --- | --- |
| **Acronym** | **Greek Region** | **N** | **Voucher** | **Latitude** | **Longitude** |
| ***Alkanna sieberi*** | | | | | |
| AT28 | Central Crete | 11 | GR-1-BBGK-19,668 | 34.92904 | 24.77959 |
| AT27 | Western Crete | 11 | GR-1-BBGK-19,667 | 35.18339 | 24.24294 |
| ***Alkanna tinctoria*** | | | | | |
| AT01 | South mainland | 9 | GR-1-BBGK-18,6127 | 37.97558 | 23.62006 |
| AT10 | South mainland | 8 | GR-1-BBGK-18,6136 | 37.87581 | 23.77331 |
| AT02 | South mainland | 9 | GR-1-BBGK-18,6128 | 37.96651 | 23.77741 |
| AT06 | South mainland | 9 | GR-1-BBGK-18,6133 | 38.03168 | 23.49035 |
| AT07 | South mainland | 9 | GR-1-BBGK-18,6134 | 37.94776 | 22.97474 |
| AT08 | South mainland | 9 | GR-1-BBGK-18,6134 | 37.91818 | 22.9967 |
| AT17 | Central mainland | 9 | GR-1-BBGK-19,509 | 39.35058 | 22.97063 |
| AT19 | Central mainland | 9 | GR-1-BBGK-19,658 | 38.94382 | 22.86016 |
| AT21 | Central mainland | 9 | GR-1-BBGK-19,660 | 38.79034 | 22.44321 |
| AT25 | Central mainland | 9 | GR-1-BBGK-19,665 | 38.50414 | 23.06436 |
| AT03 | North mainland | 11 | GR-1-BBGK-18,6081 | 40.63138 | 22.97166 |
| AT04 | North mainland | 9 | GR-1-BBGK-18,6091 | 40.64277 | 22.99777 |
| AT13 | North mainland | 8 | N/A | 40.64666 | 22.98777 |
| ATA8 | North mainland | 9 | GR-1-BBGK-18,6100 | 40.11294 | 23.31472 |

| **Table S2**. Pairwise *F*_ST_ among sampling localities of *Alkanna tinctoria* and *A. sieberi*. *F*_ST_ was estimated from 16,107 SNPs derived from RAD-seq data. | | | | | | | | | | | | | | | | |
| --- | --- | --- | --- | --- | --- | --- | --- | --- | --- | --- | --- | --- | --- | --- | --- | --- |
|  | ***A. sieberi*** | | ***A. tinctoria*** | | | | | | | | | | | | | |
|  | Crete | | Southern Greece | | | | | | Central Greece | | | | | Northern Greece | | |
|  | AT28 | AT27 | AT01 | AT10 | AT02 | AT06 | AT07 | AT08 | AT17 | AT19 | AT21 | AT25 | AT03 | | AT04 | AT13 |
| AT28 |  |  |  |  |  |  |  |  |  |  |  |  |  | |  |  |
| AT27 | 0.333 |  |  |  |  |  |  |  |  |  |  |  |  | |  |  |
| AT01 | 0.315 | 0.178 |  |  |  |  |  |  |  |  |  |  |  | |  |  |
| AT10 | 0.342 | 0.174 | 0.025 |  |  |  |  |  |  |  |  |  |  | |  |  |
| AT02 | 0.341 | 0.198 | 0.029 | 0.035 |  |  |  |  |  |  |  |  |  | |  |  |
| AT06 | 0.332 | 0.180 | 0.017 | 0.032 | 0.032 |  |  |  |  |  |  |  |  | |  |  |
| AT07 | 0.328 | 0.182 | 0.052 | 0.074 | 0.066 | 0.050 |  |  |  |  |  |  |  | |  |  |
| AT08 | 0.348 | 0.199 | 0.075 | 0.089 | 0.095 | 0.073 | 0.032 |  |  |  |  |  |  | |  |  |
| AT17 | 0.361 | 0.229 | 0.097 | 0.114 | 0.118 | 0.098 | 0.120 | 0.123 |  |  |  |  |  | |  |  |
| AT19 | 0.373 | 0.233 | 0.102 | 0.116 | 0.124 | 0.104 | 0.123 | 0.126 | 0.062 |  |  |  |  | |  |  |
| AT21 | 0.386 | 0.252 | 0.130 | 0.144 | 0.155 | 0.132 | 0.148 | 0.149 | 0.092 | 0.083 |  |  |  | |  |  |
| AT25 | 0.409 | 0.279 | 0.152 | 0.172 | 0.174 | 0.151 | 0.175 | 0.181 | 0.144 | 0.150 | 0.174 |  |  | |  |  |
| AT03 | 0.309 | 0.240 | 0.150 | 0.180 | 0.156 | 0.161 | 0.146 | 0.186 | 0.189 | 0.213 | 0.226 | 0.255 |  | |  |  |
| AT04 | 0.310 | 0.238 | 0.142 | 0.173 | 0.152 | 0.155 | 0.143 | 0.176 | 0.174 | 0.198 | 0.212 | 0.242 | 0.010 | |  |  |
| AT13 | 0.325 | 0.257 | 0.152 | 0.189 | 0.160 | 0.167 | 0.150 | 0.193 | 0.199 | 0.223 | 0.238 | 0.269 | 0.007 | | 0.010 |  |
| ATA8 | 0.338 | 0.268 | 0.162 | 0.198 | 0.166 | 0.171 | 0.163 | 0.205 | 0.210 | 0.232 | 0.251 | 0.280 | 0.071 | | 0.077 | 0.070 |

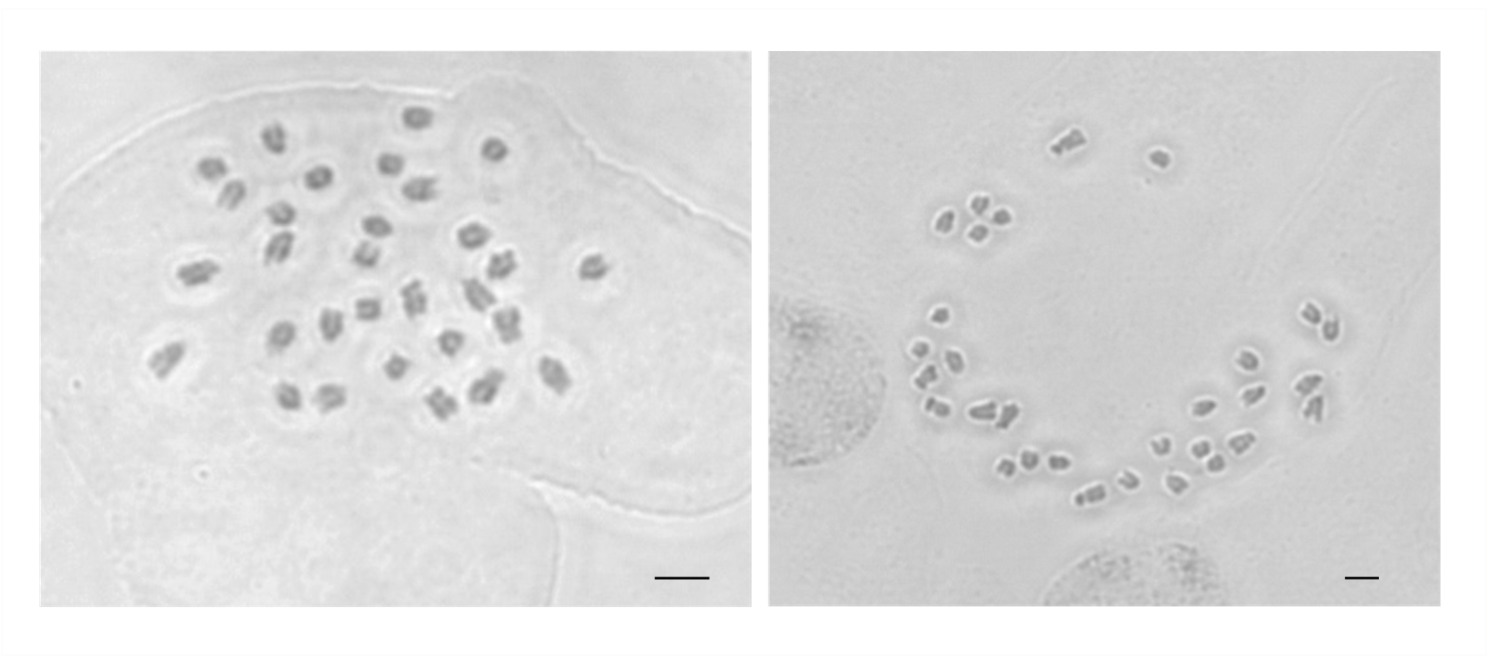
**Figure S1.** Chromosomes of *Alkanna tinctoria* (2*n* = 30). Scale bar 5µm.

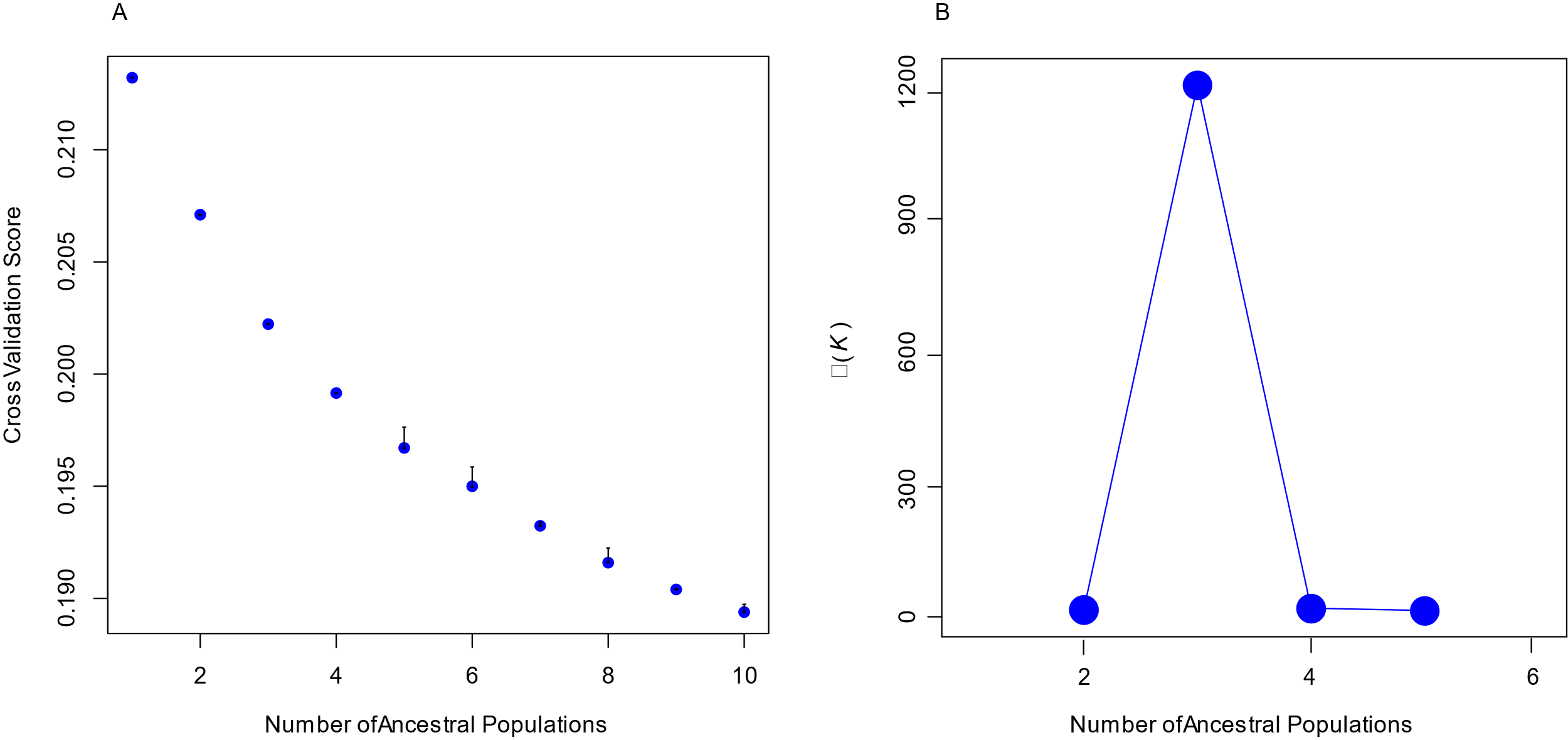

**Figure S2.** Cross validation score (A) and ΔK (B) showing optimal K identified by TESS3 and STRUCTURE HARVESTER, respectively. In the cross validation plot, the largest steps can be seen between K1 and K3. After K3, the values of cross validation score decrease at a slower rate.

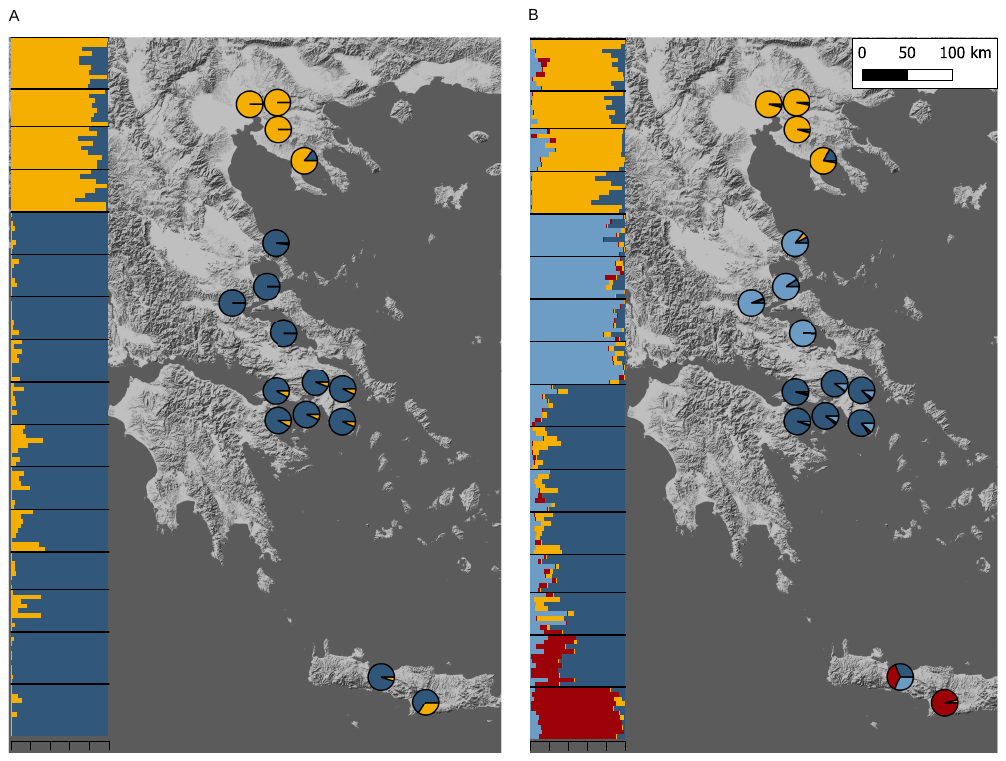

**Figure S3.** Genetic structure in studied *Alkanna* populations inferred based on EBG-derived tetraploid genotypes (n = 148). Ancestry proportions inferred with TESS3 averaged for sampling locality are plotted as pie charts. Inset shows ancestry proportions from STRUCTURE as vertical bars where each vertical bar represents an individual. Each color represents a genetic cluster. Ancestry proportions based on *K4*.

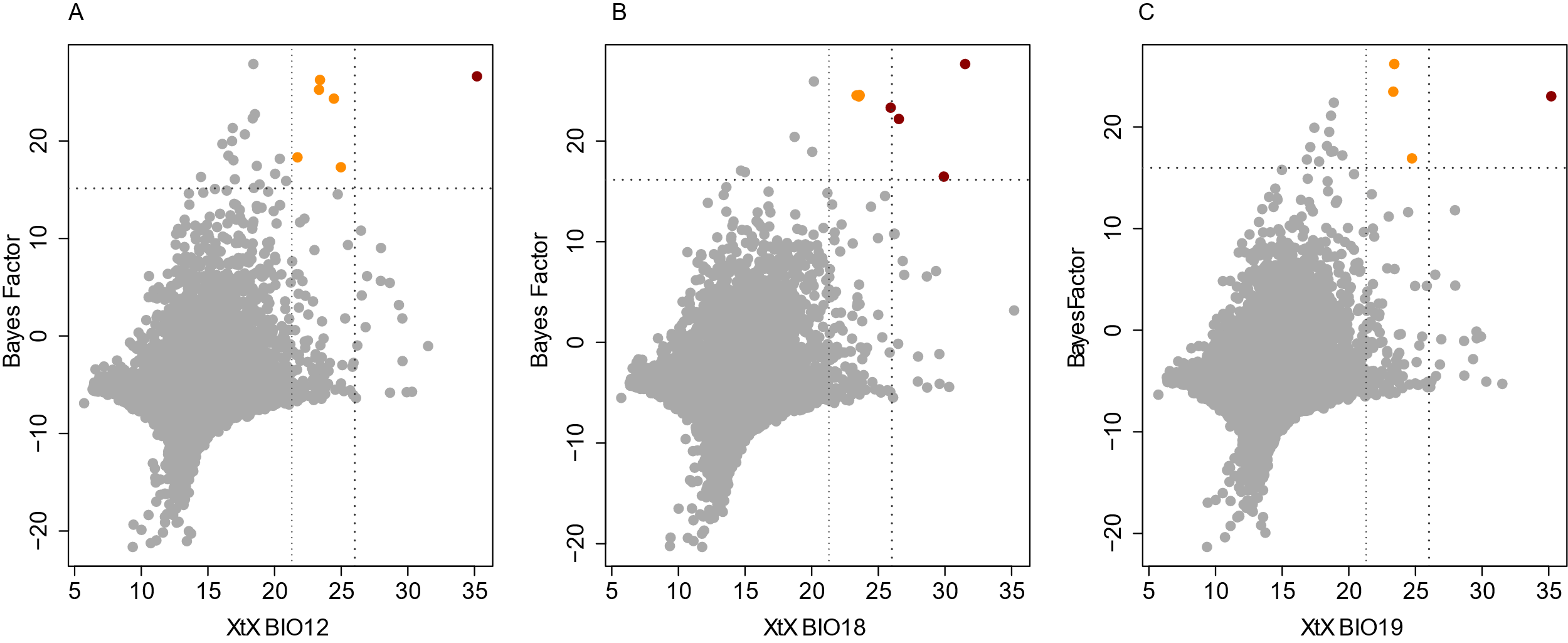

**Figure S4.** Outlier analysis of genotype-environment association (BayPass) of *Alkanna tinctoria***.** Scatter plots of correlation of XtX and Bayes factor (BF) for SNPs showing association to environmental variables. (A) BIO12, Annual Precipitation (B) BIO18, Precipitation of Warmest Quarter and (C) BIO19, Precipitation of Coldest Quarter. Scatter dots highlighted in different colors represent associated SNPs at different thresholds. Red with XtX and BF > 99.9%; orange with XtX > 99% and BF > 99.9%.

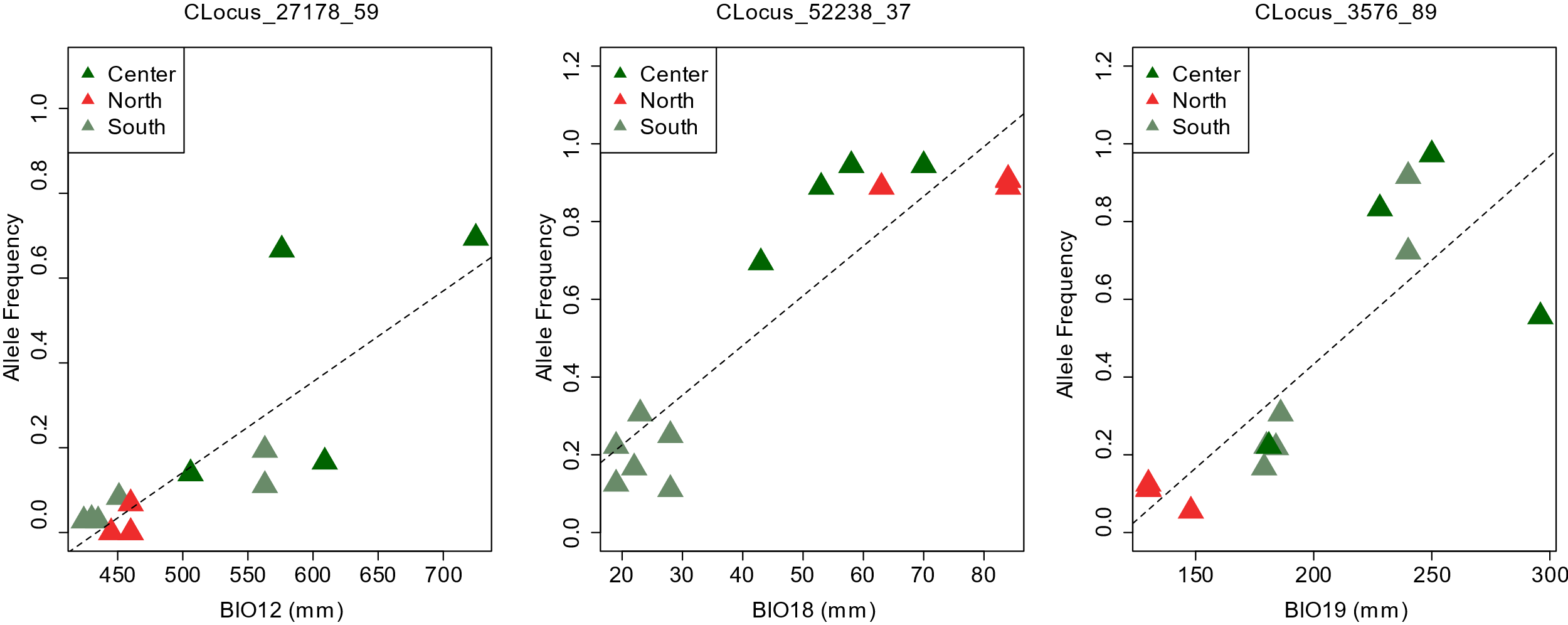

**Figure S5:** Changes in allele frequencies of loci identified by BayPass along the environmental gradient (BIO12 = Annual Precipitation, BIO18 = Precipitation of Warmest Quarter and BIO19 = Precipitation of Coldest Quarter).

**Notes S1**

Custom function used to calculate pi from EBG outputfile

##function

pi_calculator <- function(df, ploidy, total_sites){ #total length is number of loci* bp per locus

df[df == -9] <- NA

j <- rowSums(df, na.rm = T)

n <- ((ncol(df)-rowSums(is.na(df)))*ploidy)

a <- 2*j*(n-j)

b <- n*(n-1)

local_pi <- (a/b)

sum_local_pi <- sum(local_pi)

total_pi <- sum_local_pi/total_sites

return(total_pi)

}

###import your data of EBG output with rows as individuals and columns as counts of alternate #alleles

#example of run

poly <- data.frame(read.table("input.txt",sep="\t", header=T))

total_sites = 2446

ploidy = 4

###subsetting the data for populations

> AT28 = poly[,2:12]

#calculating pie

> pi_calculator(AT28, 4, 2446*94)

0.007107616
